# HHV-6B, HHV-7, and B19V Are Frequently Found DNA Viruses in the Human Thymus but Show No Definitive Link with Myasthenia Gravis

**DOI:** 10.1101/2024.06.27.600940

**Authors:** Kirsten Nowlan, Leo Hannolainen, Irini M Assimakopoulou, Pia Dürnsteiner, Joona Sarkkinen, Santeri Suokas, Lea Hedman, Pentti J. Tienari, Klaus Hedman, Mikael Niku, Leena-Maija Aaltonen, Antti Huuskanen, Jari Räsänen, Ilkka K Ilonen, Mikko I. Mäyränpää, Johannes Dunkel, Sini M Laakso, Maria Söderlund-Venermo, Maria F. Perdomo, Eliisa Kekäläinen

**Affiliations:** Translational Immunology Research Program, University of Helsinki, Helsinki, Finland; Department of Virology, University of Helsinki and Helsinki University Hospital, Helsinki, Finland; Department of Bacteriology and Immunology, University of Helsinki, Helsinki, Finland; Veterinary Biosciences, Faculty of Veterinary Medicine, University of Helsinki, Helsinki, Finland; Department of Neurology, Neurocenter, Helsinki University Hospital, Helsinki, Finland; Department of Otorhinolaryngology, Head and Neck Surgery, Helsinki University Hospital, Helsinki, Finland; Pediatric Cardiac and Transplantation Surgery Department, Helsinki Children’s Hospital, University of Helsinki, Helsinki, Finland; Department of General Thoracic and Esophageal Surgery, Heart and Lung Center, Helsinki University Hospital and University of Helsinki, Helsinki, Finland; Department of Pathology, University of Helsinki, Helsinki, Finland; HUS Diagnostic Center, HUSLAB, Clinical Microbiology, Helsinki, Finland

**Keywords:** Myasthenia gravis, Thymus, Thymoma, DNA viruses, Parvovirus B19, Herpesviruses

## Abstract

Myasthenia gravis (MG) is an autoimmune disorder characterised by autoantibodies that target components of the neuromuscular junction, primarily the acetylcholine receptor (AChR), resulting in muscle weakness. The thymus plays a significant role in MG pathogenesis, particularly in patients under the age of 50, who display pathological alterations and possess elements conducive to autoimmune reactions. Although viral infections are suspected drivers of thymic pathogenesis, the exact aetiology of MG remains elusive. This study investigates the potential link between MG and DNA viruses within the thymus. Using targeted next-generation sequencing and quantitative PCR, we analysed the presence of human parvovirus B19 (B19V) and nine human herpesviruses (HSV-1, HSV-2, VZV, EBV, CMV, HHV-6A, HHV-6B, HHV-7, and HHV-8) in fresh tissue samples from 19 non-thymomatous MG patients, 16 thymomas (3 with and 14 without MG), 41 normal thymus tissues, and 20 tonsils from healthy individuals. HHV-6B was the most common virus, found in over 50% of all tissue groups. B19V DNA was detected in 40% of adult control thymic tissue, 72% of MG thymus, 7.7% of non-MG thymoma, and 50% of tonsil samples. HHV-7 was present in 15-30% of thymus tissues and 95% of tonsils, while EBV was detected in less than 25% of all thymus samples but 85% of tonsils. In B19V seropositive individuals, B19V DNA was detected in 100% of thymic tissue from both MG patients and healthy individuals, except in thymomatous tissues, where it was found in only one of thirteen seropositive individuals. Immunohistochemistry for B19V protein expression did not show evident B19V VP1/VP2 protein expression, indicating dormant viral persistence. Laser capture microdissection (LCM) and RNAscope in situ hybridisation pinpointed B19V DNA localisation to the thymus medulla. This study is the first to demonstrate the persistence of various DNA viruses in the human thymus. However, neither B19V nor the nine human herpesviruses showed specific enrichment in MG thymic tissue compared to controls, suggesting that these viral infections are unlikely to be sole environmental triggers for MG.

## 1. Introduction

Myasthenia Gravis (MG) is a rare autoimmune disease, that is characterised by the production of autoantibodies that target the nicotinic acetylcholine receptors (AChRs) or functionally related proteins (MuSK and LRP4) at the postsynaptic membrane of the neuromuscular junction [1, 2]. The autoantibodies disrupt neuromuscular transmission, causing pronounced skeletal muscle weakness in affected individuals [2]. MG is heterogeneous both in its clinical presentation and in pathophysiology, with symptoms ranging from focal to generalised weakness, and disease onset can span from childhood to late adulthood, with peaks observed in younger women and older men [1]. This has led to the categorisation of several clinical subgroups, distinguished by autoantibody profiles, clinical features, and thymic involvement. The main subtypes of generalised MG include early-onset MG (EOMG), characterised by thymic hyperplasia and symptom onset before age 50; late-onset MG (LOMG), with less association to thymic pathologies and symptoms that emerge after age 50; and thymoma-associated MG (TAMG), featuring neoplastic thymic alterations [1, 3]. Thymectomy is most beneficial as a therapeutic treatment for myasthenia gravis in EOMG patients, particularly AChR-positive females with thymic hyperplasia. While some patients with LOMG and TAMG also respond to thymectomy, the therapeutic success in these groups is less pronounced compared to the EOMG subgroup [4, 5]. In both EOMG and TAMG, ectopic germinal centres (eGCs) often emerge within the pathologic thymus due to chronic inflammation, leading to the production of AChR autoantibodies [6, 7, 8]. While the significance of circulating antibodies against AChRs in MG pathogenesis is well-established [2, 7], the underlying aetiology of MG largely remains elusive.

Viral infections have long been recognised as potential environmental triggers for autoimmunity. Current research suggests several mechanisms through which viruses can disrupt immune tolerance and induce autoantibody production, including molecular mimicry, epitope spreading, and bystander activation [9, 10]. Notably, a strong link has been established between Epstein-Barr virus (EBV) and multiple sclerosis (MS) [11], a chronic inflammatory demyelinating disease of the central nervous system. This association suggests that EBV may play a significant role in the development of MS [11], highlighting the relevance of viruses as triggers of autoimmunity. Moreover, mounting evidence of chronic inflammation and toll-like receptor activation within the thymus of MG patients supports the hypothesis that persistent viral infection of the thymus could trigger autoimmunisation in genetically predisposed individuals [12]. Several viruses, including EBV and cytomegalovirus (CMV), have been investigated in the context of MG, however, definitive evidence of their involvement remains elusive [9, 13]. Although other herpesviruses, such as Human Herpesvirus 6 (HHV-6) and Human Herpesvirus 7 (HHV-7) have not been studied specifically in relation to MG, these viruses are known to infect CD4+ T cells and have been demonstrated to replicate within the thymus [14], making them particularly interesting in the context of the pathogenic MG thymus and deserving of further investigation.

More recently, parvovirus B19 (B19V), a single-stranded DNA virus with a narrow tropism for erythroid cells, has been associated with thymic hyperplasia in MG, suggesting it could be a potential driver of the disease [15]. Additionally, it has also been speculated that B19V infection is a plausible contributor to the formation of eGCs in TAMG [16]. However, establishing a direct link between viral infections and MG has proven challenging, mainly due to restricted access to fresh thymus material and a scarcity of mechanistic studies.

Following primary infection, B19V DNA has been shown to persist in numerous other human tissues, including tonsils, synovia, skin, kidneys, heart, brain, and more [17, 18, 19, 20]. In these non-erythroid tissues, the infection typically remains dormant (i.e. no infectious virus is produced), yet the virus may still have the potential to induce pathological effects through indirect mechanisms, such as triggering inflammatory and autoimmune processes [21, 22]. Given that B19V has been shown to persist in B cells in tonsil and intestinal lymphoid tissues [17, 22] it is plausible that the presence of B19V within the thymus of patients with MG can act as a trigger or driver for the chronic inflammation and thymic hyperplasia leading to autoantibody production. Our objective was to comprehensively assess the potential relationship between DNA viruses and thymic pathologies associated with the development of MG. We initially screened samples using NGS and followed up with qPCR on a larger cohort for B19V and nine herpesviruses (HSV-1, HSV-2, VZV, EBV, CMV, HHV-6A, HHV-6B, HHV-7 and HHV-8). This approach aimed to unveil the significance of viral infections or the persistence of DNA viruses within the MG thymus and their potential role in the aetiology and pathogenesis of MG.

## 2. Materials and Methods

### 2.1 Study Subjects and Samples

A conveniency cohort of pathologic thymic tissue samples were obtained from MG and non-MG patients undergoing therapeutic thymectomy at the Helsinki University Hospital Heart and Lung Center between 2020-2023. After gross evaluation by a pathologist (HUS Diagnostic Center Pathology), a portion of the sample, not required for clinical diagnostics, was used for the study. MG diagnoses were made in the Department of Neurology at Helsinki University Hospital and both the MG and histopathological diagnoses were confirmed through patient records from Helsinki University Hospital. To establish comparative control groups, we obtained thymic tissue samples from paediatric patients who were immunologically healthy and undergoing cardiac surgery at the New Children’s Hospital, Helsinki University Hospital (HUS/124/2023 § 19/2023). We also acquired adult thymus biopsies from individuals undergoing thoracic surgery (ERB approval HUS/151/2022 § 37/2022) and tonsil tissue samples from patients with no history of autoimmune disorders (ERB approval HUS/462/2021). Tissue samples were immediately processed as three distinct preparations. The first tissue sample was placed in an Eppendorf tube and stored at -80°C. A second larger section was embedded in OCT compound, rapidly frozen with dry ice, and stored at -80°C. The final portion was fixed in 10% formalin and embedded in paraffin. Blood samples were collected from all patients during surgery, along with samples from age- and sex-matched controls for MG patients. Plasma was separated by centrifugation and stored in aliquots at - 80°C. All study subjects or their legal guardians gave a written informed consent prior to sampling.

### 2.2 Detection of DNA Viruses using NGS

DNA extracts were mechanically fragmented using a Covaris E220 instrument, with target size of 200 bp. The library preparation was performed using the KAPA Hyperplus kit (Roche) according to the manufacturer’s protocol using unique double index adapters (Roche). For viral DNA capture, a custom panel of biotinylated RNA oligonucleotides (myBaits Custom DNA-Seq, Arbor Biosciences) was used [23]. Each individual sample underwent two rounds of hybridization, adhering to the manufacturer’s recommendations for low-input DNA (MyBaits v5 kit Arbor Biosciences) and using KAPA Universal Enhancing Oligos ((Roche) to prevent non-specific binding. These RNA-probes had a length of 100 bp and were designed complementary to the complete sequences of 42 human DNA viruses with 2X tiling. Included in the design were parvovirus B19, cutavirus, torque teno viruses 1, 10, and 13, nine herpesviruses, polyomaviruses 1–13, hepatitis B virus, papillomavirus types 2, 6, 11, 16, 18, 21, 45, bocaviruses 1–4, simian virus 40, and variola viruses minor and major. The libraries were quantified using the KAPA Library Quantification Kit (Roche) and pooled for sequencing in NovaSeq 6000 (one lane, S4, PE151, Illumina). The libraries underwent 3 × 13–25 cycles of amplification, with clean-up using KAPA Pure Beads (Roche). Negative controls (PCR-grade water) were included in library preparation, enrichment, and sequencing.

### 2.3 NGS data analysis

The data analysis was done with TRACESPipeLite, a streamlined version of TRACESPipe [24]. The paired-end reads were trimmed and collapsed with AdapterRemoval, cutting ambiguous bases at the 5’/3’ termini with quality scores below or equal to two. Reads shorter than 20 bases were discarded. FALCON-meta [25] was used to find the highest similar reference from the NCBI viral database. The reads were aligned with BWA [26] using a seed length of 1000 and a maximum diff of 0.01. Read duplicates were removed with SAMtools [27] and the consensus sequences reconstructed with BCFtools [28]. The coverage profiles were created with BEDtools [29]. When in low breadth coverage (< 15 %), the individual reads were manually inspected and confirmed by BLAST. The pipeline is freely available, under MIT licence, at https://github.com/viromelab/TRACESPipeLite, along with the code (included in the TRACESPipeLite repository).

### 2.4 Detection of DNA Viruses by qPCR

The DNA extraction was performed with the QIAamp DNA Mini Kit (Qiagen), following the manufacturer’s protocol for tissue extraction, except for the addition of a digestion step with collagenase and an increased volume of proteinase K [17]. In brief, the tissue samples were dissected into smaller sections with sterile scalpels and incubated in 100 μl of PBS and 50 μl of Liberase TL (1 mg/ml; Roche) for one hour at +37°C. After the incubation, 180 μl of buffer ATL and 40 μl of proteinase K were added and incubated at +56°C shaking for 3 hours. To detect and quantify B19V DNA, a pan-B19V qPCR assay targeting the NS1 genes of all three genotypes was employed, as previously described [30]. For the nine human herpesviruses, a three-tube multiplex qPCR assay (HSV-1, HSV-2, VZV, EBV, CMV, HHV-6A, HHV-6B, HHV-7 and HHV-8) was applied to detect each virus [31]. The amplification of B19V and herpesvirus DNA was performed with the AriaMx Real-Time PCR System (Agilent). The human gene *RNase P* was quantified both as an internal control and to normalise the viral copy numbers per million cells across all the samples [30]. The qPCR reagents, sample handling, and DNA extractions, were each performed in separate hoods and rooms to avoid contamination.

### 2.5 B19V Serology

Serum samples were analysed for acute and past infections of parvovirus B19V by IgM and IgG enzyme immunoassays (EIAs), with biotinylated virus-like particles (VLP) of VP2 as antigens [32]. Epitope-type specific (ETS) IgG towards linear peptide and conformational VLP antigens was also conducted to confirm prior immunity [33, 34].

### 2.6 Laser Capture Microdissection

We included three thymic samples from patients with EOMG that exhibited eGCs in the thymus and selected three tonsil samples with the highest B19V DNA viral loads for laser capture microdissection. Zeiss MembraneSlide 1.0 PEN were heat-treated at 180°C for 4 hours and exposed to UV light for 30 minutes at a distance of 400 mm before mounting the tissue sections. Serial sections from the OCT tissue blocks were cut with a cryostat at -20°C, each 12-μm thick, and dried on the membrane slides. Immediately the slides were stained with toluidine blue as described by Bahreini et al., 2022 [35]. B-cell follicles from thymi, and areas of the cortex and medulla, were dissected using laser microdissection with the PALM Microbeam (Carl Zeiss MicroImaging GmbH, Bernried, Germany) according to the manufacturer’s instructions. The dissected tissues from each region of each sample were collected into the cap of a 0.5-mL Zeiss Opaque AdhesiveCap tubes and stored at -80°C. To further process the samples, the same methods as earlier for the genomic DNA extraction and qPCR were used.

### 2.7 Immunohistochemistry

Immunohistochemical staining was performed according to the manufacturer’s instructions on formalin-fixed paraffin-embedded (FFPE) sections of 3-µm-thick thymus and tonsil tissues using monoclonal antibodies against B19V VP1/VP2 (R92F6, Novocastra, Newcastle, UK). Protein staining was considered positive if granular brown reaction products could be observed in the nucleus. Placental tissue from B19V-infected individuals served as a positive control and PCR-negative tissue samples as negative controls [22]. All IHC stainings were performed at an accredited clinical pathology laboratory (HUS Diagnostic Center, Pathology).

### 2.8 RNAscope *in situ* hybridization

To identify the persistence sites of B19V, RNAscope ISH (RISH) technology (Advanced Cell Diagnostics [ACD], Newark, CA) was used to detect viral DNA/RNA in 5-μm-thick FFPE thymus and tonsil tissue sections, mounted on glass slides (SuperFrost Plus). Viral nucleic acid detection was employed using 20 pairs of double-Z oligonucleotide probes designed to target the sense B19V *NS1* gene (Probe-V-B19-NS1, 967-2,215 bp; GenBank: NC_000883.2), as previously described [22]. The hybridised probe was amplified using RNAscope 2.5 HD Reagent Kit-RED (ACD) according to the manufacturer’s protocol (adjustment: Protease Plus, 7 minutes incubation), presenting the targets as red dots. Probes targeting the human housekeeping gene *PPIB* and the bacterial gene *dapB* (ACD) served as positive and negative technical controls, respectively, and B19V PCR-negative tonsil and thymus, as negative virus controls.

### 2.9 Statistical Analysis

Chi-square test was applied for comparison of seroprevalence between individuals with MG and controls of similar ages. To express viral frequency as a proportion and facilitate the comparison of the relative distribution of different virus types within each group, normalisation was performed. Fisher’s-exact test was employed for analysis of the viral frequency proportions between groups. Kruskal-Wallis test, followed by the post hoc Dunn’s test was used to measure the significance between B19V DNA viral copy numbers in the tissues from different groups of seropositive individuals. A p-value threshold of 0.05 was set to determine significant statistical comparisons. All analysis was done in in RStudio (Version 2024.04.1+748)

## 3. Results

### 3.1 Detection of DNA Viruses by NGS and quantitative PCR

DNA was extracted from fresh-frozen thymic tissue samples from patients with MG, individuals with non-MG related thymoma, and controls without thymic pathology. Additionally, DNA was extracted from palatine-tonsil tissue of adult controls. These DNA samples underwent an initial screening for the detection of viral pathogens (Figure 1A). This screening was conducted through viral genome sequencing using targeted next-generation sequencing (NGS), which employed a custom panel specifically designed for DNA viruses [23]. Based on an initial pilot study, conducted on 16 samples, B19V, HHV-6B, and HHV-7 were identified in multiple samples (Supplementary Table 2). Subsequently, qPCR for Pan-B19V, and HERQ9 multiplex qPCR for herpesviruses, were used to validate the NGS findings. Moreover, the qPCR was further utilised to assess the persistence of these viruses within the tissue from an expanded cohort (Figure 1B), which included a broader range of controls such as palatine tonsils and paediatric thymus samples. B19V DNA was most commonly found in LOMG samples (12/13), which had no thymic pathologies and featured typical involution, whereas the B19V-DNA prevalences were lower in EOMG samples (5/7), adult controls (4/11), and tonsil tissues (10/20) (Supplementary Table 3). Overall, B19V was more commonly detected in MG tissues compared to those of the controls, but this difference did not reach statistical significance (Figure 1B). However, it is essential to account for the age distributions within these groups, as the likelihood of a primary infection increases during childhood. In line with this, B19V DNA was infrequently detected in paediatric thymic samples (1/30), aligning with a seroprevalence of 2/30 children. Notably, the seropositive paediatric patient with the negative tissue finding for B19V DNA exhibited an acute infection, rather than a past infection. Surprisingly, B19V DNA was also rarely detected in both TAMG and non-MG thymoma tissues (1/16), despite the advanced age and high seroprevalence in these individuals (13/16). HHV-6B DNA exhibited a steady genoprevalence across all examined thymic tissue groups, ranging from 53% to 70%, while showing a notably higher prevalence (90%) in tonsil tissue samples. Likewise, HHV-7 DNA was identified at slightly lower but consistent prevalence within thymus tissues, ranging from 15% to 36% across various thymic subsets, but found in 95% of tonsils. EBV DNA was found to be sparsely present in thymic tissue, detected in only 5/22 MG samples (including 2 from EOMG patients), non-MG 3/14 thymoma samples, and 4/41 healthy thymus tissue samples, and at low copy numbers (Figure 1B). In contrast, EBV was more frequently identified in tonsils, where 17/20 individuals tested positive and displayed higher copy numbers per one million cells. Regarding CMV, its DNA was detected in 2/22 MG samples and in 3/54 thymus tissue controls (healthy and thymoma samples), while it was absent in all tonsil tissue samples (Figure 1B).

**Figure 1.**
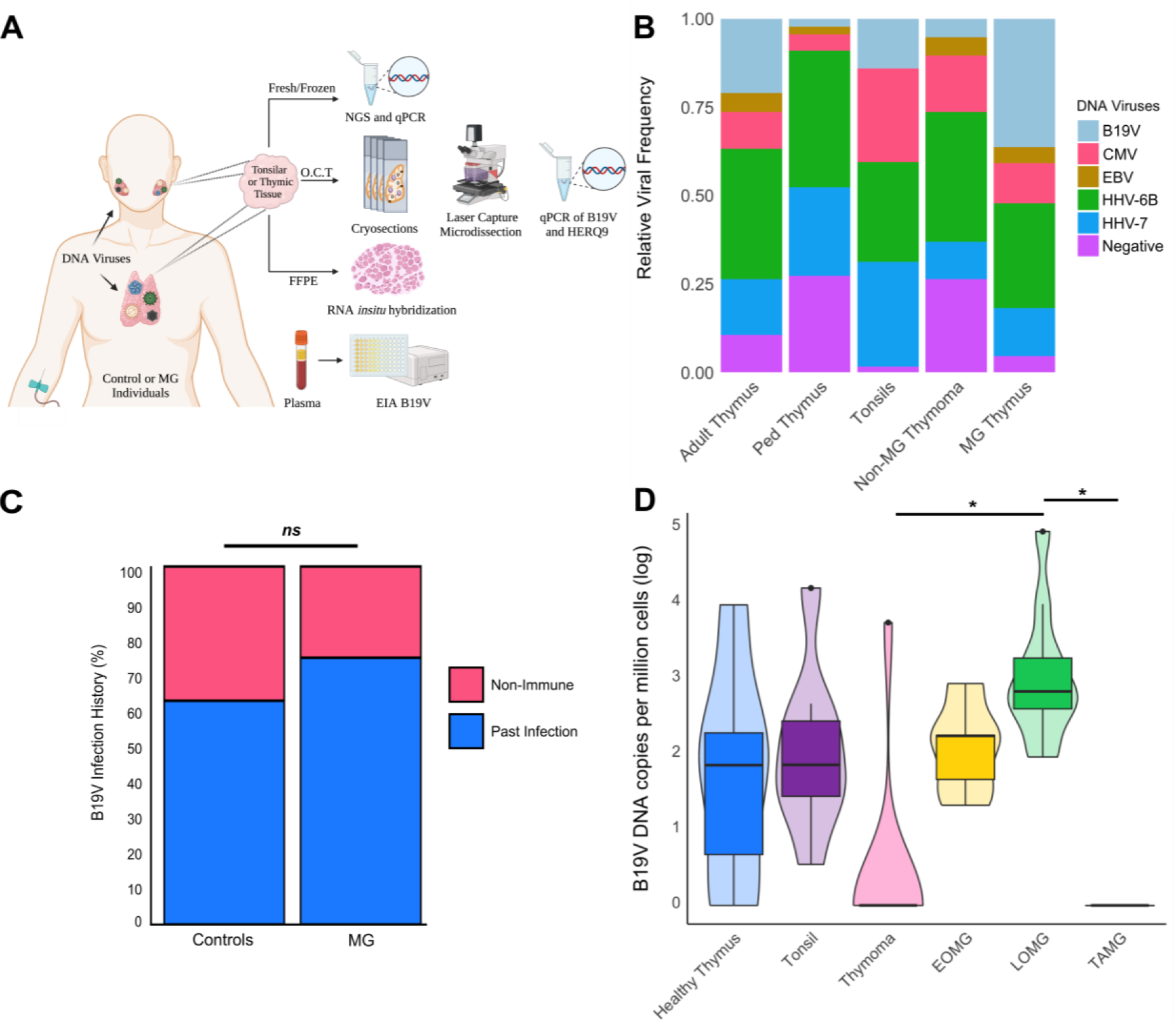
Comparative analysis of DNA-virus presence in MG thymic tissues and controls, accompanied by serological screening for B19V in paired blood samples. **A)** Schematic representation of the study design. Created with BioRender.com **B)** Stacked bar plot illustrating the relative frequency of viral detection in each tissue group. The ’Negative’ group represents tissues PCR-Negative for all screened DNA viruses (herpesviruses and B19V). No significant difference in virus frequency was detected between the various tissue groups (p =1). **C)** Serological evaluation of B19V seroprevalence utilising paired plasma samples from the original MG cohort and controls, supplemented by an archival cohort of plasma from MG patients and controls. Paediatric samples were excluded from this analysis due to age bias. The median age ± SD of the MG group was 36±18.3 years (n=51), whereas for the control group, it was 42.5±19 years (n=56). No significant difference in the past B19V infection history between MG patients and age-matched controls could be detected (p =0.26). **D)** Box plot illustrating B19V copy numbers per one million cells (normalised for *RNase* P) in the different subsets of MG compared to control tissues. Significant differences in viral loads could be detected between the LOMG group and the thymoma and TAMG groups (p =0.0002, p =0.0105, respectively).

### 3.2 Serological Testing for B19V and Normalised B19V Copy Numbers in Tissues of Seropositive Individuals

Serological testing was conducted on plasma obtained from paired blood samples to assess B19V infection history. Our analysis revealed no significant difference, p = 0.26, in the seroprevalence between MG patients and age- and sex-matched controls (Figure 1C) (Table 1). Here we were also able to utilise archival plasma samples from a sizable cohort of EOMG patients in addition to the original cohort, where a substantial subset of MG patients that exhibited thymic pathologies such as hyperplasia (8/23) and ectopic germinal centres (5/23), were seronegative for B19V (Supplementary Table 1).

**Table 1.** Cohort Characteristics

|  | B19V Tissue qPCR |  |  |  | B19V Serology |  |  |  |
| --- | --- | --- | --- | --- | --- | --- | --- | --- |
| Sample Type | N | Median Age (IQR) | Male (%) | Female (%) | N | Median Age (IQR) | Male (%) | Female (%) |
| **MG Patients** | **22** | **54.5 (39.5-67.8)** | **54.5** | **45.5** | **51** | **36.0 (26.0-53.5)** | **31.4** | **68.6** |
| EOMG | 7 | 27.0 (19.5-33.5) | 14.3 | 85.7 | 33 | 28.0 (26.0-53.5) | 9.1 | 90.9 |
| LOMG | 12 | 67.5 (57.0-70.5) | 75.0 | 25.0 | 15 | 65.0 (55.5-69.0) | 73.3 | 26.7 |
| TAMG | 3 | 49.0 (47.5-50.5) | 66.7 | 33.3 | 3 | 49.0 (47.5-50.5) | 66.7 | 33.3 |
| **Controls** | **74** | **21.5 (5.1-42.2)** | **40.5** | **59.5** | **77** | **26.0 (8.6-56.0)** | **44.2** | **55.8** |
| Paediatric Thymus | 30 | 4.0 (2.1-6.0) | 53.3 | 46.7 | 21 | 5.0 (3.6-6.0) | 57.1 | 42.9 |
| Adult Thymus | 11 | 52.0 (34.5-68.0) | 45.5 | 54.5 | 24 | 52.5 (39.8-63.8) | 50.0 | 50.0 |
| Non-MG Thymoma | 13 | 60.0 (56.0-70.0) | 46.2 | 53.8 | 13 | 58.0 (55.0-69.0) | 53.8 | 46.2 |
| Tonsils | 20 | 25.0 (21.0-26.2) | 15.0 | 85.0 | 19 | 25.0 (21.0-26.0) | 15.8 | 84.2 |

Among seropositive individuals with available fresh tissues, qPCR confirmed viral DNA presence in all cases except for both TAMG and non-MG thymoma tissue samples. Comparative analysis of viral copy numbers among seropositive individuals revealed no discernible difference between EOMG patients and any of the control groups. However, LOMG samples exhibited significantly higher B19V DNA copy numbers per million cells compared to both the TAMG and thymoma tissue samples, which largely tested negative via qPCR despite being from seropositive individuals.

### 3.3 Adipose tissue in the thymus as a reservoir of B19V DNA

Notably, LOMG had the highest positivity rate of B19V DNA (Supplementary Figure 1), and upon histological examination, the predominant tissue composition in LOMG samples was adipose tissue, suggesting a potential reservoir for B19V persistence. In a subset of thymoma samples, we were able to dissect adjacent adipose tissue and separately digest it for qPCR. In these cases, adipose tissue from seropositive individuals consistently yielded positive results for B19V DNA, while the corresponding thymoma tissue from the same individual tested negative by qPCR (Table 2). This observation further supports the idea that adipose tissue could be a potential reservoir for B19V persistence.

**Table 2:** B19V qPCR results from thymoma tissue and adjacent adipose tissue. *BDL= below detection limit*

| Sample | Sample Type | Rnase P<br>(copies/ul) | B19V per<br>million cells | HHV-6B per<br>million cells | HHV-7 per<br>million cells |
| --- | --- | --- | --- | --- | --- |
| Thymus 45 | Thymoma<br>Tissue | 7.51e+05 | BDL | BDL | BDL |
| Thymus 45 | Adjacent<br>Adipose<br>Tissue | 2.52e+04 | 1.33e+04 | BDL | BDL |
| Thymus 36 | Thymoma Tissue | 1.77e+06 | BDL | 4.68e+02 | BDL |
| Thymus 36 | Adjacent Adipose Tissue | 4.37e+04 | 2.19e+02 | BDL | BDL |
| Thymus 28 | Thymoma Tissue | 1.62e+05 | BDL | 9.94e+01 | BDL |
| Thymus 28 | Adjacent Adipose Tissue | 9.35e+04 | 3.49e+01 | 4.02e+01 | 1.70e+01 |

### 3.4 Detecting B19V VP1/VP2 Proteins with Immunohistochemistry

To understand whether the persistent B19V infection might manifest as a productive tissue infection, as outlined previously [15], we employed immunohistochemistry (IHC) to identify active production of B19V VP1/VP2 proteins. We detected faint nonspecific signals that displayed a non-nuclear staining pattern and were found across all tissues, irrespective of seropositivity or seronegativity. No distinctly positive cells expressing B19V VP1/VP2 were evident in any thymic tissues when compared to a positive control sample. Likewise, the tonsil tissue exhibited one or two darker stains that could also be observed in the PCR-negative control tonsil tissue (Figure 2), indicating they are nonspecific signals. In all, our results are consistent with persisting B19V DNA in the thymus but without production of viral structural proteins.

**Figure 2.**
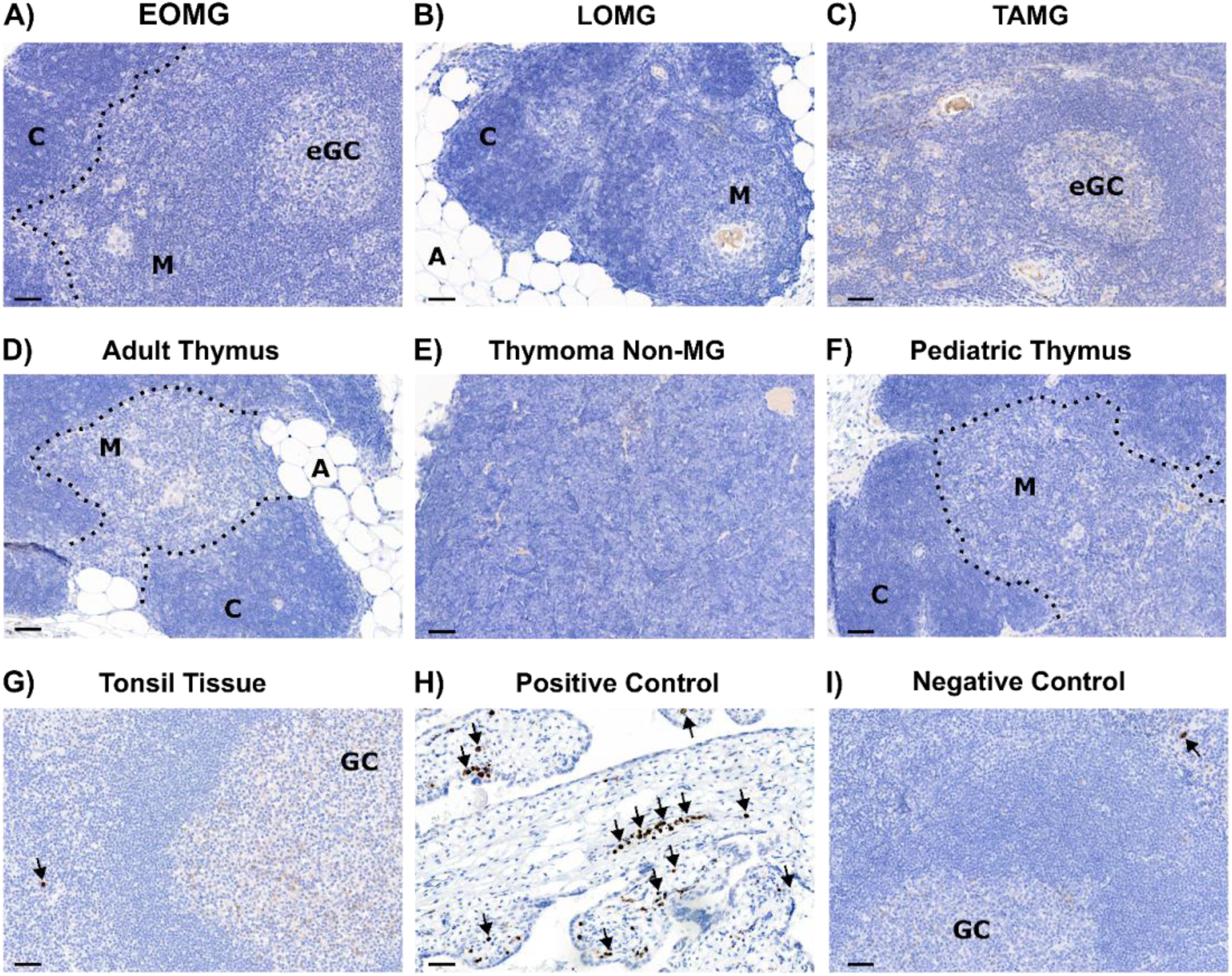
Immunohistochemical Staining of B19V VP1/VP2 in Various Subsets of MG Thymic Tissues and Control Tissues. A-F) B19V PCR-positive thymus tissue from seropositive individuals. A) Hyperplastic thymic tissue from an EOMG patient with eGCs present within the tissue. B) Involuted thymic tissue from a LOMG patient, in which thymic islets within adipose tissue are evident. C) Thymoma tissue from a patient with TAMG and eGCs present within the tissue. D) Healthy thymic tissue from an adult control sample which was macroscopically involuted, i.e thymic islets surrounded by adipose tissue. E) Thymoma tissue from a patient that does not have MG. F) Healthy thymic tissue from a paediatric individual, with little to no involution evident. B19V VP1/VP2 antigen staining was negative in all thymic tissues. G) B19V PCR-positive palatine tonsil tissue from an immunologically healthy B19V seropositive individual and genopositive. A dark nuclear signal could be detected in a few cells within the whole tissue section. H) Positive control placental tissue from an individual with an active B19V infection. Dark nuclear signals can be seen in multiple cells within the tissue. I) B19V PCR-negative control palatine tonsil tissue from a seronegative individual; sparse dark nuclear signals can also be seen in a pattern similar to that of the qPCR-positive tonsil, indicating that this signal is likely to be nonspecific. Tissue images are 20X and the scale bar is 50µm. M = medulla, C = cortex, A= adipose, eGC = ectopic germinal centre, GC= germinal centre.

### 3.5 Persistent B19V DNA is located in the thymic medulla but not found in the autoantibody-producing eGCs

To explore the microanatomical niche where B19V and herpesviruses may persist within the thymus, we utilised Laser Capture Microdissection (LCM) to isolate the ectopic germinal centres from frozen tissue samples of EOMG patients, alongside regions of the medulla and cortex. Despite the challenge posed by low viral copy numbers of B19V detected from whole tissue samples, analysis of LCM-isolated tissue from the same samples with the highest B19V DNA levels revealed that B19V DNA is predominantly present in the thymic medulla, with no detectable concentrations in the eGCs or cortex. Notably, HHV-6B was detected in both eGCs and medullary regions of these samples. Adequate DNA yields were confirmed in each sample, as evidenced by the *RNase* P values. To corroborate the B19V observation, we employed RNAscope in situ hybridization (RISH), which revealed positive signals exclusively within the medullary region of the thymus. In a PCR-negative EOMG sample from a seronegative individual, no signal was detected. Interestingly, however, in tonsil tissue, B19V DNA was detected primarily in the germinal centres (Figure 3).

**Figure 3.**
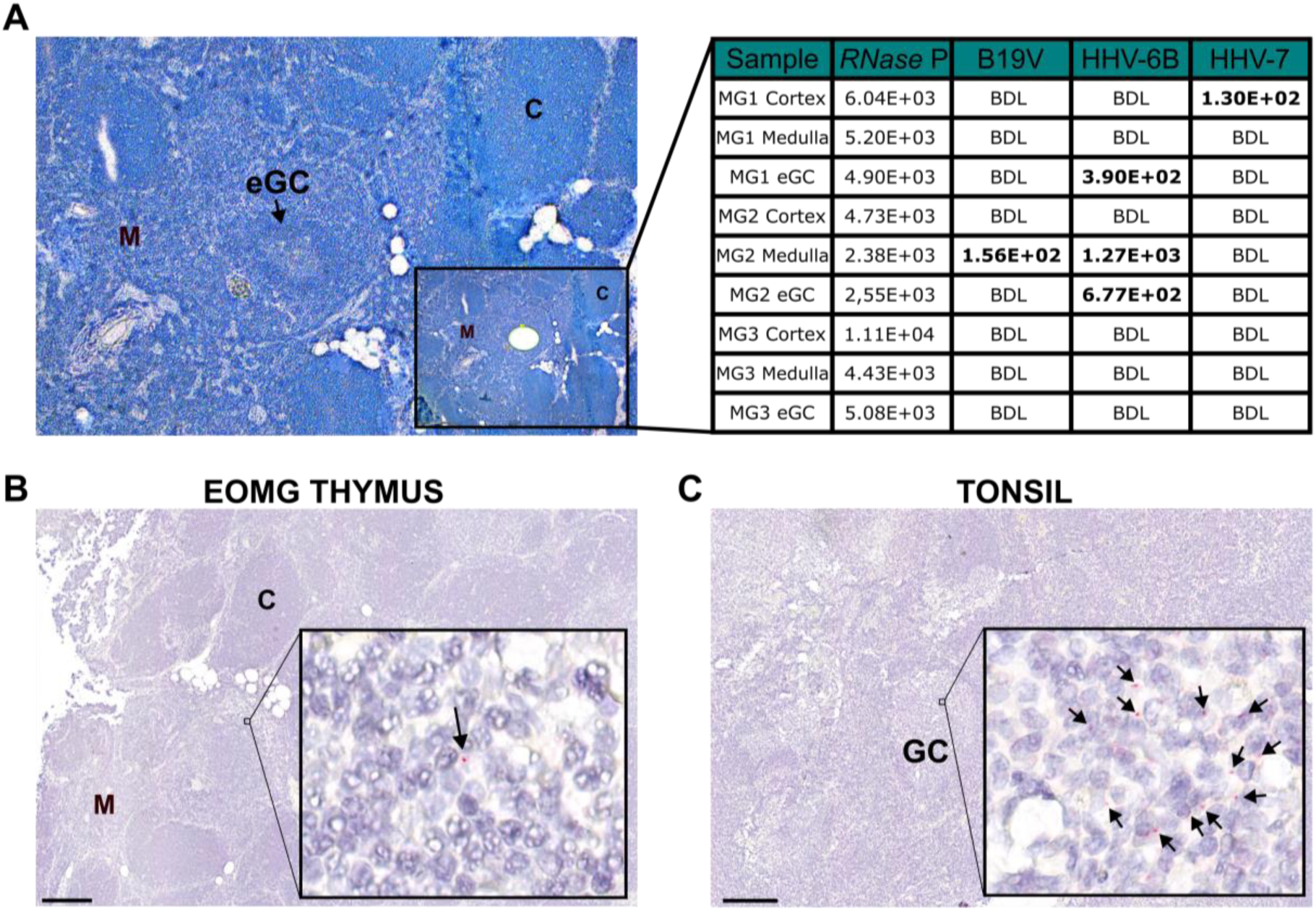
Laser Capture Microdissection and qPCR from regions of EOMG thymi, and RNAscope in situ hybridization (RISH) of EOMG thymus and tonsil tissue from a control. **A)** Cryosections of EOMG thymi were stained with Toluidine Blue, followed by laser capture microdissection to isolate distinct regions: ectopic germinal centres (eGCs), medulla, and cortical areas, each collected into separate tubes (n=3). Subsequently, qPCR analysis targeting all herpesviruses and B19V was performed. RN*ase* P values confirmed adequate tissue collection across all regions and samples. Notably, B19V-DNA detection was limited to the medullary region with the highest B19V copy numbers per one million cells (from whole tissue), whereas HHV-6B DNA was detected in both the medulla and ectopic germinal centres (eGCs) of the tissue, and HHV-7 DNA in the cortex of a single sample. BDL= Below Detection Limit **B)** RISH for B19V nucleic acids on EOMG thymic tissue revealed sparse DNA positive signals distributed throughout the tissue but localised within the medullary regions devoid of eGCs. **C)** B19V RISH on PCR-positive tonsillar control tissues demonstrated DNA positive signals primarily within germinal centres. Scale Bar = 200µm, M = medulla, C = cortex, eGC = ectopic germinal centre, GC= germinal centre.

## 4. Discussion

The role of the thymus in the pathogenesis of MG has been extensively researched [1, 3, 36]. However, identifying the precise molecular trigger that initiates and fosters the development of thymic follicular hyperplasia-related MG has remained challenging. Viral infection or reactivation has been proposed as a potential trigger for MG, as viral-related inflammatory responses may induce chronic inflammation, instigate autoimmune responses or modify the host transcriptome [9, 13, 37]. Utilising a comprehensive methodology with fresh tissues from unique material of controls and newly diagnosed MG patients, as well as paired blood samples from these individuals, we expansively screened an extensive array of DNA viruses using a large targeted NGS panel. This then further directed in-depth analysis using multiplexed qPCR, to investigate whether specific DNA viruses are directly linked to the etiopathogenesis of MG in individuals with thymic pathologies. However, our investigation indicates no direct causal relationship between a specific DNA virus within the thymus and the pathological alterations contributing to MG. Nevertheless, our comprehensive study does demonstrate the persistence of multiple DNA viruses, notably HHV-6B, HHV-7, and B19V, within the thymus. This persistence is observed across various groups, including samples with normal thymic histology and no MG, as well as in samples where the individual was afflicted with MG regardless of thymic pathology, with the exception of B19V in thymoma tissue of individuals both with and without MG (Figure 1B and 1D). While not specific to MG, our findings underscore the susceptibility of the thymus to viral infection and persistence, with potential implications for thymic function that warrant further exploration.

Previous studies have indicated the presence of B19V proteins within the thymus of individuals with MG, suggesting a potentially productive infection within the pathogenic tissue [15]. However, there is insufficient evidence supporting the notion that B19V could replicate within non-erythroid cells or tissues [38, 39, 40]. It has been shown that B19V can enter nonpermissive cells, such as immune and endothelial cells, with the help of antibody-dependent enhancement (ADE), via the Fc receptor or complement factor C1q [17, 38, 41]. Additionally, while B19V DNA has been detected in various tissue types, the potential for B19V reactivation in previously healthy individuals remains uncertain, as conclusive evidence for such an occurrence has yet to be demonstrated [42]. To address these uncertainties, we conducted IHC for B19V VP1/VP2 within our cohort, aiming to replicate previous findings suggesting a productive infection [15, 16]. However, we were unable to detect specific viral protein in any tissue other than the positive control; placental tissue from a woman with an active B19V infection (Figure 2). Therefore, we conclude that the persistence of B19V within the thymus seems to be dormant, with no viral expression or subsequent replication, either in individuals with MG or in the thymus or tonsil controls. Nevertheless, RT-PCR to determine if viral mRNA was produced, could not be performed as the tissue was not stored in an RNA preservative.

Upon analysing the B19V serostatus of individuals in our sample collection, we observed persistence of the viral DNA within thymic tissue in all previously infected individuals, except in cases of thymoma tissue. The B19V seroprevalence rate showed no significant difference between patients with MG and controls, aligning with the reported seroprevalence of B19V in pregnant women from Finland of similar age to most cases of EOMG diagnosis, approximately 60% [43, 44]. These findings contrast with the relationship between EBV and MS, where complete seropositivity of EBV was observed in large cohorts of MS patients [11, 45]. In our cohort, the seroprevalence data does not support any causal link between B19V and MG. Age matching between controls and disease groups is crucial due to the increased likelihood of exposure and infection in higher age groups. Our study included controls within the corresponding age brackets due to the bimodal distribution of MG incidence (Table 1) [1, 46]. Additionally, there were no significant differences in B19V DNA copy numbers between the different groups, except for LOMG and thymoma tissues. These results lead us to conclude that B19V is unlikely to play a causative role in the onset of thymic pathologies or MG. Moreover, our observation of a lack of B19V in thymoma tissue, irrespective of MG status, indicates that thymoma tissue is a less conducive environment for B19V persistence compared to normal thymic tissue. This presents an intriguing avenue for further research into the mechanisms underlying B19V persistence within non-erythroid cells or tissues, as well as the factors making thymoma a less conducive environment for B19V persistence.

HHV-6, known for its tropism for T cells, presents a potential area for exploration, particularly regarding a potential reactivation in the human thymus. While this remains an understudied area, a study of primary infection in mice showed thymic depletion, highlighting the novel nature of our findings [47, 48]. Here we describe HHV-6B and HHV-7 as commonly persisting viruses within the thymus, of not only individuals with MG but also that of healthy controls. It is imperative to understand how viral persistence in the thymus may impact its microenvironment, potentially impairing the process of T-cell generation and thymic function. This is of particular relevance in the context of haematopoietic stem cell transplantation (HSCT) where HHV-6 viremia has been shown to affect T-cell reconstitution [49].

Interestingly, EBV and CMV are rarely detected within the thymus, which contrasts with their prevalence in the tonsils of young adults. While previous literature comprises various studies, including some focusing solely on the serological prevalence and others on retrospective analysis of thymic tissues, there remains controversy regarding the involvement of EBV in the onset of MG [50, 51]. However, our findings do not support the notion that EBV is “enriched” or even persistent within the thymus of MG, or specifically EOMG, patients. The absence of EBV detection within the thymus of EOMG patients raises intriguing questions. Despite patients with eGCs experiencing an influx of B cells, the tissue remains EBV-DNA negative, contrary to what would be expected based on statistical seroprevalence. This raises the possibility that eGCs could originate from thymic B cells rather than peripheral B cells, suggesting a potential protective mechanism of the thymus against viral infection [52].

Formalin fixation, although effective for morphological and molecular preservation, can induce DNA damage and cross-linking, hindering nucleic acid screening, especially for low-abundance targets like persistent DNA viruses [53]. Hence, our rigorous approach of using fresh-frozen tissue and paired blood samples (to evaluate infection history) allowed for a more comprehensive and sensitive analysis of DNA viruses within thymic tissue, despite a limited sample size due to the rarity of MG. While no direct causative DNA virus was identified, this does not entirely rule out their contribution to MG aetiology. Bystander activation of viral infections or the presence of multiple permissive viruses in the thymus could lead to chronic inflammation [42, 54, 55]. It should be noted that our focus was solely on DNA viruses, and further exploration of RNA viruses is warranted. Mechanistic studies, such as challenging thymic tissue with infectious agents, could provide insights into the inflammatory pathways triggered by viral infections. Additionally, investigating other viruses will elucidate the role of viral infection in the pathogenesis of MG. This underscores the need for future research in this understudied area to understand the implications of viral persistence on thymic function and MG development.

## Acknowledgments

We are grateful to the patients and healthy controls for their participation; without them, this study would not have been possible. We thank Marjo Rissanen for her help with sample processing. Additionally, we extend our thanks to Mikko Räsänen, and Eva Sutinen, for their contribution to patient procurement. The authors would also like to thank the Genome Biology Unit, supported by HiLIFE and the Faculty of Medicine, University of Helsinki, and Biocenter Finland, for their imaging services.

## Funding

JPRD JTC2019 and Research Council of Finland (308913) (EK)

Doctoral Programme in Integrative Life Science (ILS), University of Helsinki (KN)

Finnish Medical Foundation (MFP),

Life and Health Medical Support Association (MFP, MSV),

The Sigrid Jusélius Foundation (MSV)

Helsinki University Hospital Research and Education Fund (KH),

Finska Läkaresällskapet (KH, MFP),

Magnus Ehrnrooth foundation (KH),

Jane and Aatos Erkko Foundation (KH)

## Author contributions

Conceptualization: KN, LH, IMA, MSV, MFP, EK

Methodology: KN, LH, IMA, PD, MIM, LH, KH

Validation: KN, LH, IMA, EK, MSV, MFP

Formal Analysis: KN, LH, IMA

Investigation: KN, LH, IMA, PD, JS, IKI, LH, SS, MIM, JD, SML, MFP

Resources: MIM, IKI, KN, SS, SML, MFP, LH, JR, AH, PT, LMA, JD, MSV, EK

Data Curation: KN, LH, JS

Writing – Original Draft: KN, LH, IMA

Writing – Review & Editing: KN, LH, IMA, JS, MIM, IKI, MN, SS, SML, MFP, KH, JR, AH, PT, MSV, EK

Visualization: KN

Supervision: MIM, SML, KH, MSV, MFP, EK

Project administration: KN, MFP, EK

Funding acquisition: MIM, KH, MSV, MFP, EK

## Supplements

**Supplementary Table 1:** Patient characteristics from extended cohort of individuals with MG

| Sample | Age Group (y) | Sex | MG Type | B19V Serostatus | eGCs | Hyperplasia |
| --- | --- | --- | --- | --- | --- | --- |
| MGO1 | 15-30 | F | EOMG | Non-Immune | Yes | Yes |
| MGO2 | 15-30 | F | EOMG | Non-Immune | Yes | Yes |
| MG03 | 15-30 | F | EOMG | Non-Immune | Unknown | Unknown |
| MG04 | 15-30 | F | EOMG | Non-Immune | Yes | Yes |
| MG05 | 15-30 | F | EOMG | Non-Immune | Yes | Yes |
| MG06 | 30-50 | F | EOMG | Non-Immune | Unknown | Unknown |
| MG07 | 30-50 | F | EOMG | Non-Immune | Yes | Yes |
| MG08 | 30-50 | F | EOMG | Non-Immune | No | Yes |
| MG09 | >50 | M | LOMG | Non-Immune | Unknown | Unknown |
| MG10 | >50 | F | LOMG | Non-Immune | Yes | No |
| MG11 | 15-30 | F | EOMG | Past B19V<br>infection | Unknown | Unknown |
| MG12 | 15-30 | F | EOMG | Past B19V<br>infection | Unknown | Unknown |
| MG13 | 15-30 | F | EOMG | Past B19V<br>infection | Yes | Yes |
| MG14 | 15-30 | F | EOMG | Past B19V<br>infection | Yes | Yes |
| MG15 | 15-30 | F | EOMG | Past B19V<br>infection | Unknown | Unknown |
| MG16 | 15-30 | F | EOMG | Past B19V<br>infection | Yes | Yes |
| MG17 | 15-30 | F | EOMG | Past B19V<br>infection | Yes | Yes |
| MG18 | 15-30 | F | EOMG | Past B19V<br>infection | Yes | Yes |
| MG19 | 15-30 | F | EOMG | Past B19V<br>infection | No | Yes |
| MG20 | 30-50 | F | EOMG | Past B19V<br>infection | Yes | Yes |
| MG21 | 30-50 | F | EOMG | Past B19V<br>infection | Unknown | Unknown |
| MG22 | 30-50 | F | EOMG | Past B19V<br>infection | Yes | Yes |
| MG23 | 30-50 | F | EOMG | Past B19V<br>infection | Yes | Yes |
| MG24 | 30-50 | F | EOMG | Past B19V<br>infection | No | Yes |
| MG25 | 30-50 | F | EOMG | Past B19V<br>infection | Unknown | Unknown |
| MG26 | 30-50 | F | EOMG | Past B19V<br>infection | Yes | Yes |
| MG27 | 30-50 | M | EOMG | Past B19V<br>infection | Unknown | Unknown |
| MG28 | 30-50 | M | EOMG | Past B19V<br>infection | Unknown | Unknown |
| MG29 | >50 | M | LOMG | Past B19V<br>infection | Unknown | Unknown |

**Supplementary Table 2:** Breadth coverage of DNA viruses for pilot study of 16 samples by targeted NGS screening.

| Sample | Age | Sex | MG | MG<br>Type | B19V | HHV-6B | HHV-7 | TTV | MCPyV |
| --- | --- | --- | --- | --- | --- | --- | --- | --- | --- |

|  | Group (y) |  |  |  |  |  |  |  |  |
| --- | --- | --- | --- | --- | --- | --- | --- | --- | --- |
| Thymus 1 | 30-50 | M | 1 | EOMG | - | - | - | - | - |
| Thymus 2 | 30-50 | M | 1 | EOMG | 93 | - | - | - | - |
| Thymus 3 | >50 | F | 0 | NA | - | - | - | - | - |
| Thymus 4 | >50 | M | 0 | NA | - | 7 | - | - | - |
| Thymus 5 | 15-30 | F | 1 | EOMG | 11 | - | - | - | - |
| Thymus 6 | >50 | F | 0 | NA | - | - | - | - | - |
| Thymus 7 | >50 | F | 1 | LOMG | 14 | - | - | - | - |
| Thymus 8 | >50 | M | 1 | LOMG | 18 | 6 | - | - | - |
| Thymus 9 | 15-30 | F | 1 | EOMG | 23 | - | - | - | - |
| Thymus 10 | >50 | M | 1 | LOMG | 99 | 14 | 13 | - | - |
| Thymus 11 | >50 | F | 0 | NA | - | - | - | - | - |
| Thymus 12 | >50 | F | 0 | NA | - | - | - | - | - |
| Thymus 13 | >50 | M | 0 | NA | 99 | 46 | 6,7 | 6,6 | - |
| Thymus 14 | 30-50 | F | 0 | NA | 94 | 7 | - | - | 3 |
| Thymus 15 | >50 | M | 1 | TAMG | 32 | - | - | - | - |
| Thymus 16 | 30-50 | F | 0 | NA | 92 | 14 | 1 | - | - |

**Supplementary Table 3:**
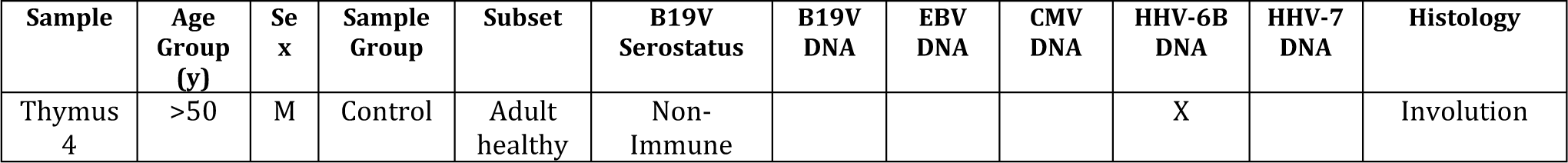
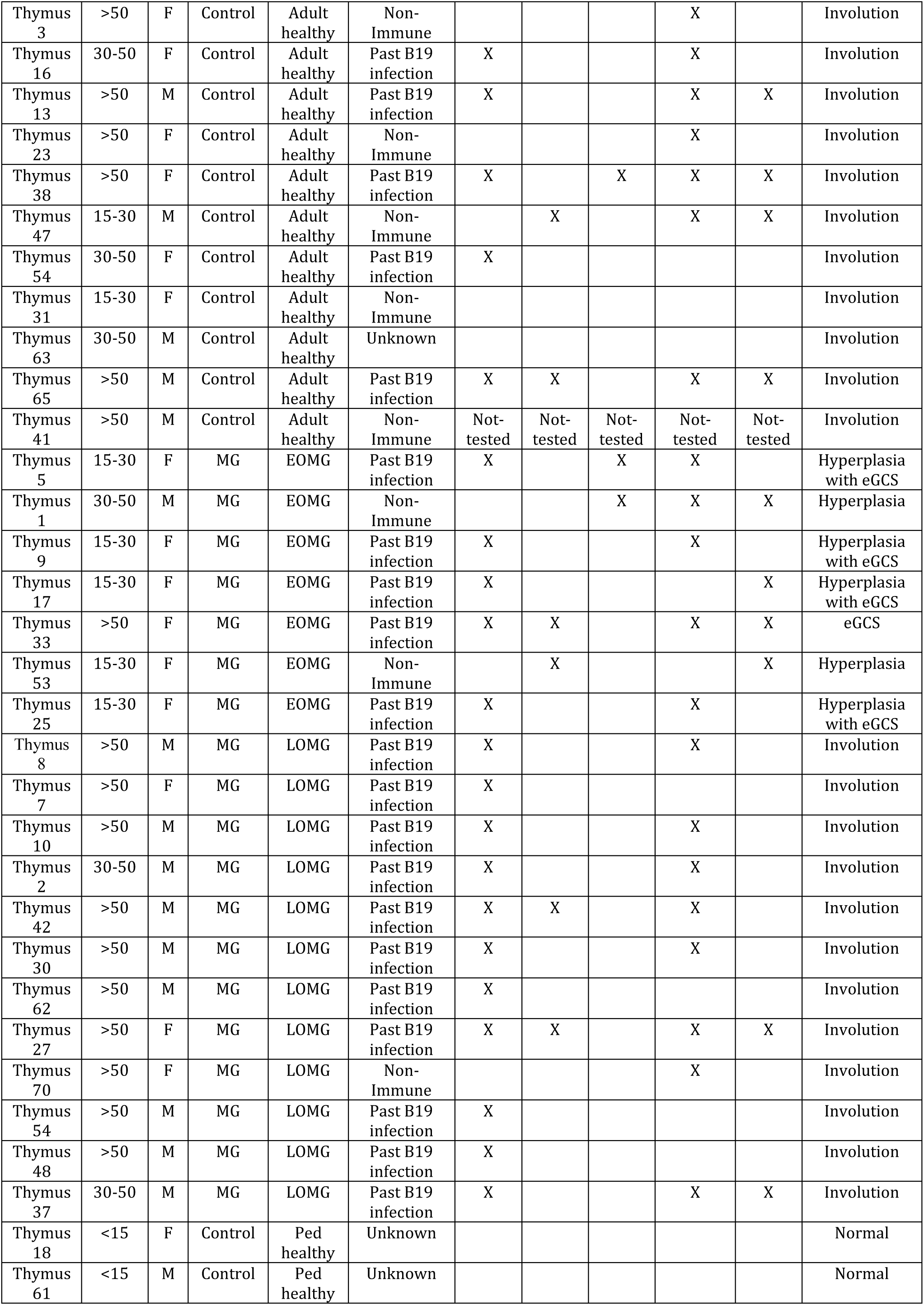
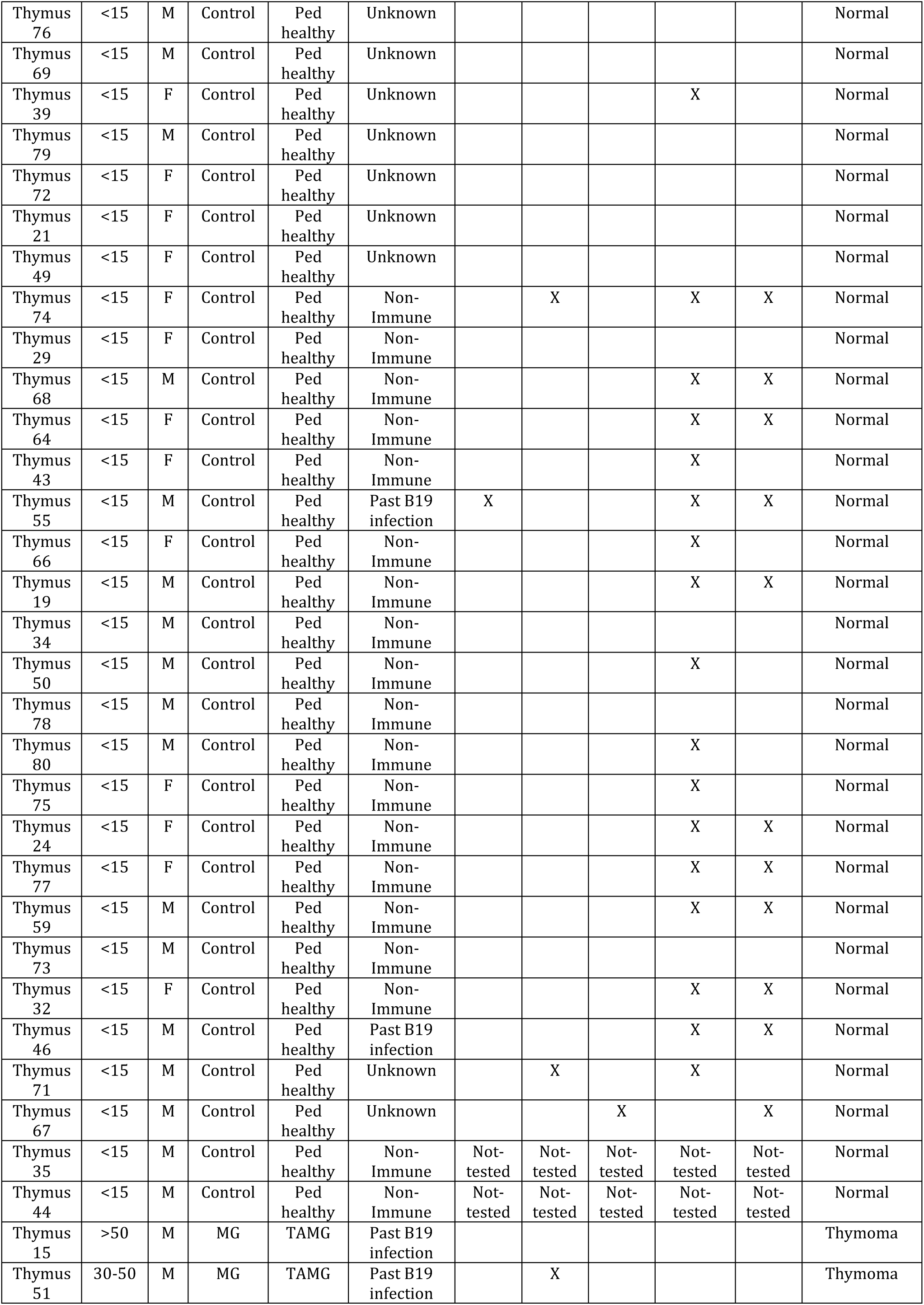
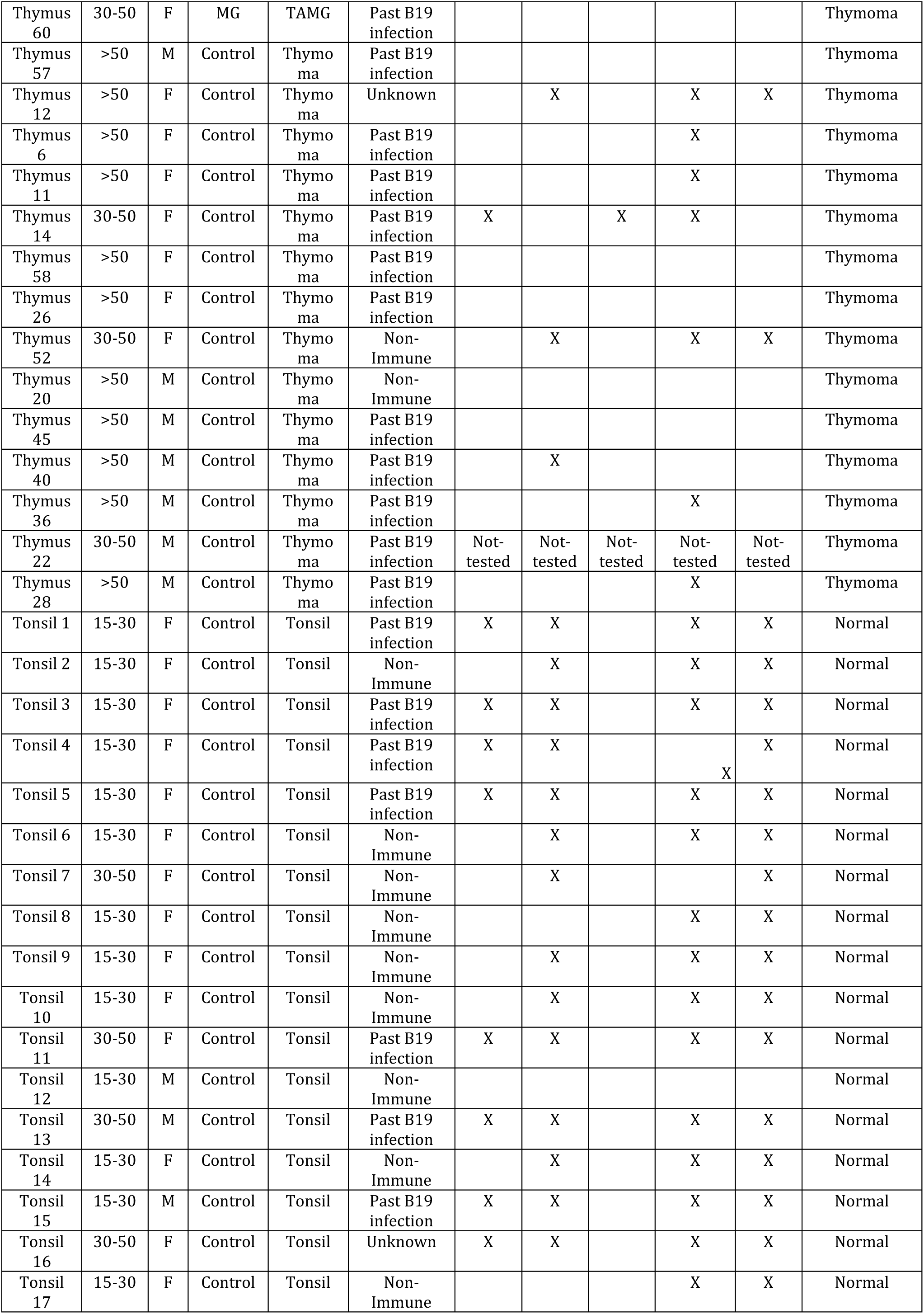
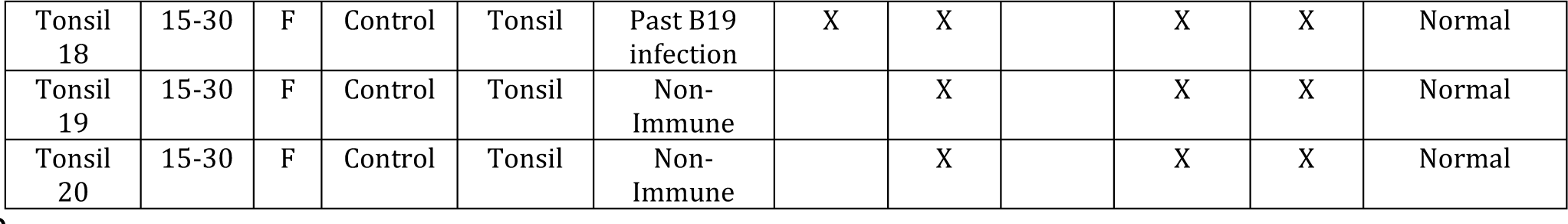
Cohort characteristics, and qPCR detection of DNA viruses from fresh tissue samples

| Sample | Age Group (y) | Sex | Sample Group | Subset | B19V Serostatus | B19V DNA | EBV DNA | CMV DNA | HHV-6B DNA | HHV-7 DNA | Histology |
| --- | --- | --- | --- | --- | --- | --- | --- | --- | --- | --- | --- |
| Thymus 4 | >50 | M | Control | Adult healthy | Non-Immune |  |  |  | X |  | Involution |
| Thymus 3 | >50 | F | Control | Adult healthy | Non-Immune |  |  |  | X |  | Involution |
| Thymus 16 | 30-50 | F | Control | Adult healthy | Past B19 infection | X |  |  | X |  | Involution |
| Thymus 13 | >50 | M | Control | Adult healthy | Past B19 infection | X |  |  | X | X | Involution |
| Thymus 23 | >50 | F | Control | Adult healthy | Non-Immune |  |  |  | X |  | Involution |
| Thymus 38 | >50 | F | Control | Adult healthy | Past B19 infection | X |  | X | X | X | Involution |
| Thymus 47 | 15-30 | M | Control | Adult healthy | Non-Immune |  | X |  | X | X | Involution |
| Thymus 54 | 30-50 | F | Control | Adult healthy | Past B19 infection | X |  |  |  |  | Involution |
| Thymus 31 | 15-30 | F | Control | Adult healthy | Non-Immune |  |  |  |  |  | Involution |
| Thymus 63 | 30-50 | M | Control | Adult healthy | Unknown |  |  |  |  |  | Involution |
| Thymus 65 | >50 | M | Control | Adult healthy | Past B19 infection | X | X |  | X | X | Involution |
| Thymus 41 | >50 | M | Control | Adult healthy | Non-Immune | Not-tested | Not-tested | Not-tested | Not-tested | Not-tested | Involution |
| Thymus 5 | 15-30 | F | MG | EOMG | Past B19 infection | X |  | X | X |  | Hyperplasia with eGCS |
| Thymus 1 | 30-50 | M | MG | EOMG | Non-Immune |  |  | X | X | X | Hyperplasia |
| Thymus 9 | 15-30 | F | MG | EOMG | Past B19 infection | X |  |  | X |  | Hyperplasia with eGCS |
| Thymus 17 | 15-30 | F | MG | EOMG | Past B19 infection | X |  |  |  | X | Hyperplasia with eGCS |
| Thymus 33 | >50 | F | MG | EOMG | Past B19 infection | X | X |  | X | X | eGCS |
| Thymus 53 | 15-30 | F | MG | EOMG | Non-Immune |  | X |  |  | X | Hyperplasia |
| Thymus 25 | 15-30 | F | MG | EOMG | Past B19 infection | X |  |  | X |  | Hyperplasia with eGCS |
| Thymus 8 | >50 | M | MG | LOMG | Past B19 infection | X |  |  | X |  | Involution |
| Thymus 7 | >50 | F | MG | LOMG | Past B19 infection | X |  |  |  |  | Involution |
| Thymus 10 | >50 | M | MG | LOMG | Past B19 infection | X |  |  | X |  | Involution |
| Thymus 2 | 30-50 | M | MG | LOMG | Past B19 infection | X |  |  | X |  | Involution |
| Thymus 42 | >50 | M | MG | LOMG | Past B19 infection | X | X |  | X |  | Involution |
| Thymus 30 | >50 | M | MG | LOMG | Past B19 infection | X |  |  | X |  | Involution |
| Thymus 62 | >50 | M | MG | LOMG | Past B19 infection | X |  |  |  |  | Involution |
| Thymus 27 | >50 | F | MG | LOMG | Past B19 infection | X | X |  | X | X | Involution |
| Thymus 70 | >50 | F | MG | LOMG | Non-Immune |  |  |  | X |  | Involution |
| Thymus 54 | >50 | M | MG | LOMG | Past B19 infection | X |  |  |  |  | Involution |
| Thymus 48 | >50 | M | MG | LOMG | Past B19 infection | X |  |  |  |  | Involution |
| Thymus 37 | 30-50 | M | MG | LOMG | Past B19 infection | X |  |  | X | X | Involution |
| Thymus 18 | <15 | F | Control | Ped healthy | Unknown |  |  |  |  |  | Normal |
| Thymus 61 | <15 | M | Control | Ped healthy | Unknown |  |  |  |  |  | Normal |
| Thymus 76 | <15 | M | Control | Ped healthy | Unknown |  |  |  |  |  | Normal |
| Thymus 69 | <15 | M | Control | Ped healthy | Unknown |  |  |  |  |  | Normal |
| Thymus 39 | <15 | F | Control | Ped healthy | Unknown |  |  |  | X |  | Normal |
| Thymus 79 | <15 | M | Control | Ped healthy | Unknown |  |  |  |  |  | Normal |
| Thymus 72 | <15 | F | Control | Ped healthy | Unknown |  |  |  |  |  | Normal |
| Thymus 21 | <15 | F | Control | Ped healthy | Unknown |  |  |  |  |  | Normal |
| Thymus 49 | <15 | F | Control | Ped healthy | Unknown |  |  |  |  |  | Normal |
| Thymus 74 | <15 | F | Control | Ped healthy | Non-Immune |  | X |  | X | X | Normal |
| Thymus 29 | <15 | F | Control | Ped healthy | Non-Immune |  |  |  |  |  | Normal |
| Thymus 68 | <15 | M | Control | Ped healthy | Non-Immune |  |  |  | X | X | Normal |
| Thymus 64 | <15 | F | Control | Ped healthy | Non-Immune |  |  |  | X | X | Normal |
| Thymus 43 | <15 | F | Control | Ped healthy | Non-Immune |  |  |  | X |  | Normal |
| Thymus 55 | <15 | M | Control | Ped healthy | Past B19 infection | X |  |  | X | X | Normal |
| Thymus 66 | <15 | F | Control | Ped healthy | Non-Immune |  |  |  | X |  | Normal |
| Thymus 19 | <15 | M | Control | Ped healthy | Non-Immune |  |  |  | X | X | Normal |
| Thymus 34 | <15 | M | Control | Ped healthy | Non-Immune |  |  |  |  |  | Normal |
| Thymus 50 | <15 | M | Control | Ped healthy | Non-Immune |  |  |  | X |  | Normal |
| Thymus 78 | <15 | M | Control | Ped healthy | Non-Immune |  |  |  |  |  | Normal |
| Thymus 80 | <15 | M | Control | Ped healthy | Non-Immune |  |  |  | X |  | Normal |
| Thymus 75 | <15 | F | Control | Ped healthy | Non-Immune |  |  |  | X |  | Normal |
| Thymus 24 | <15 | F | Control | Ped healthy | Non-Immune |  |  |  | X | X | Normal |
| Thymus 77 | <15 | F | Control | Ped healthy | Non-Immune |  |  |  | X | X | Normal |
| Thymus 59 | <15 | M | Control | Ped healthy | Non-Immune |  |  |  | X | X | Normal |
| Thymus 73 | <15 | M | Control | Ped healthy | Non-Immune |  |  |  |  |  | Normal |
| Thymus 32 | <15 | F | Control | Ped healthy | Non-Immune |  |  |  | X | X | Normal |
| Thymus 46 | <15 | M | Control | Ped healthy | Past B19 infection |  |  |  | X | X | Normal |
| Thymus 71 | <15 | M | Control | Ped healthy | Unknown |  | X |  | X |  | Normal |
| Thymus 67 | <15 | M | Control | Ped healthy | Unknown |  |  | X |  | X | Normal |
| Thymus 35 | <15 | M | Control | Ped healthy | Non-Immune | Not-tested | Not-tested | Not-tested | Not-tested | Not-tested | Normal |
| Thymus 44 | <15 | M | Control | Ped healthy | Non-Immune | Not-tested | Not-tested | Not-tested | Not-tested | Not-tested | Normal |
| Thymus 15 | >50 | M | MG | TAMG | Past B19 infection |  |  |  |  |  | Thymoma |
| Thymus 51 | 30-50 | M | MG | TAMG | Past B19 infection |  | X |  |  |  | Thymoma |
| Thymus 60 | 30-50 | F | MG | TAMG | Past B19 infection |  |  |  |  |  | Thymoma |
| Thymus 57 | >50 | M | Control | Thymoma | Past B19 infection |  |  |  |  |  | Thymoma |
| Thymus 12 | >50 | F | Control | Thymoma | Unknown |  | X |  | X | X | Thymoma |
| Thymus 6 | >50 | F | Control | Thymoma | Past B19 infection |  |  |  | X |  | Thymoma |
| Thymus 11 | >50 | F | Control | Thymoma | Past B19 infection |  |  |  | X |  | Thymoma |
| Thymus 14 | 30-50 | F | Control | Thymoma | Past B19 infection | X |  | X | X |  | Thymoma |
| Thymus 58 | >50 | F | Control | Thymoma | Past B19 infection |  |  |  |  |  | Thymoma |
| Thymus 26 | >50 | F | Control | Thymoma | Past B19 infection |  |  |  |  |  | Thymoma |
| Thymus 52 | 30-50 | F | Control | Thymoma | Non-Immune |  | X |  | X | X | Thymoma |
| Thymus 20 | >50 | M | Control | Thymoma | Non-Immune |  |  |  |  |  | Thymoma |
| Thymus 45 | >50 | M | Control | Thymoma | Past B19 infection |  |  |  |  |  | Thymoma |
| Thymus 40 | >50 | M | Control | Thymoma | Past B19 infection |  | X |  |  |  | Thymoma |
| Thymus 36 | >50 | M | Control | Thymoma | Past B19 infection |  |  |  | X |  | Thymoma |
| Thymus 22 | 30-50 | M | Control | Thymoma | Past B19 infection | Not-tested | Not-tested | Not-tested | Not-tested | Not-tested | Thymoma |
| Thymus 28 | >50 | M | Control | Thymoma | Past B19 infection |  |  |  | X |  | Thymoma |
| Tonsil 1 | 15-30 | F | Control | Tonsil | Past B19 infection | X | X |  | X | X | Normal |
| Tonsil 2 | 15-30 | F | Control | Tonsil | Non-Immune |  | X |  | X | X | Normal |
| Tonsil 3 | 15-30 | F | Control | Tonsil | Past B19 infection | X | X |  | X | X | Normal |
| Tonsil 4 | 15-30 | F | Control | Tonsil | Past B19 infection | X | X |  | X | X | Normal |
| Tonsil 5 | 15-30 | F | Control | Tonsil | Past B19 infection | X | X |  | X | X | Normal |
| Tonsil 6 | 15-30 | F | Control | Tonsil | Non-Immune |  | X |  | X | X | Normal |
| Tonsil 7 | 30-50 | F | Control | Tonsil | Non-Immune |  | X |  |  | X | Normal |
| Tonsil 8 | 15-30 | F | Control | Tonsil | Non-Immune |  |  |  | X | X | Normal |
| Tonsil 9 | 15-30 | F | Control | Tonsil | Non-Immune |  | X |  | X | X | Normal |
| Tonsil 10 | 15-30 | F | Control | Tonsil | Non-Immune |  | X |  | X | X | Normal |
| Tonsil 11 | 30-50 | F | Control | Tonsil | Past B19 infection | X | X |  | X | X | Normal |
| Tonsil 12 | 15-30 | M | Control | Tonsil | Non-Immune |  |  |  |  |  | Normal |
| Tonsil 13 | 30-50 | M | Control | Tonsil | Past B19 infection | X | X |  | X | X | Normal |
| Tonsil 14 | 15-30 | F | Control | Tonsil | Non-Immune |  | X |  | X | X | Normal |
| Tonsil 15 | 15-30 | M | Control | Tonsil | Past B19 infection | X | X |  | X | X | Normal |
| Tonsil 16 | 30-50 | F | Control | Tonsil | Unknown | X | X |  | X | X | Normal |
| Tonsil 17 | 15-30 | F | Control | Tonsil | Non-Immune |  |  |  | X | X | Normal |
| Tonsil 18 | 15-30 | F | Control | Tonsil | Past B19 infection | X | X |  | X | X | Normal |
| Tonsil 19 | 15-30 | F | Control | Tonsil | Non-Immune |  | X |  | X | X | Normal |
| Tonsil 20 | 15-30 | F | Control | Tonsil | Non-Immune |  | X |  | X | X | Normal |

**Supplementary Figure 1:**
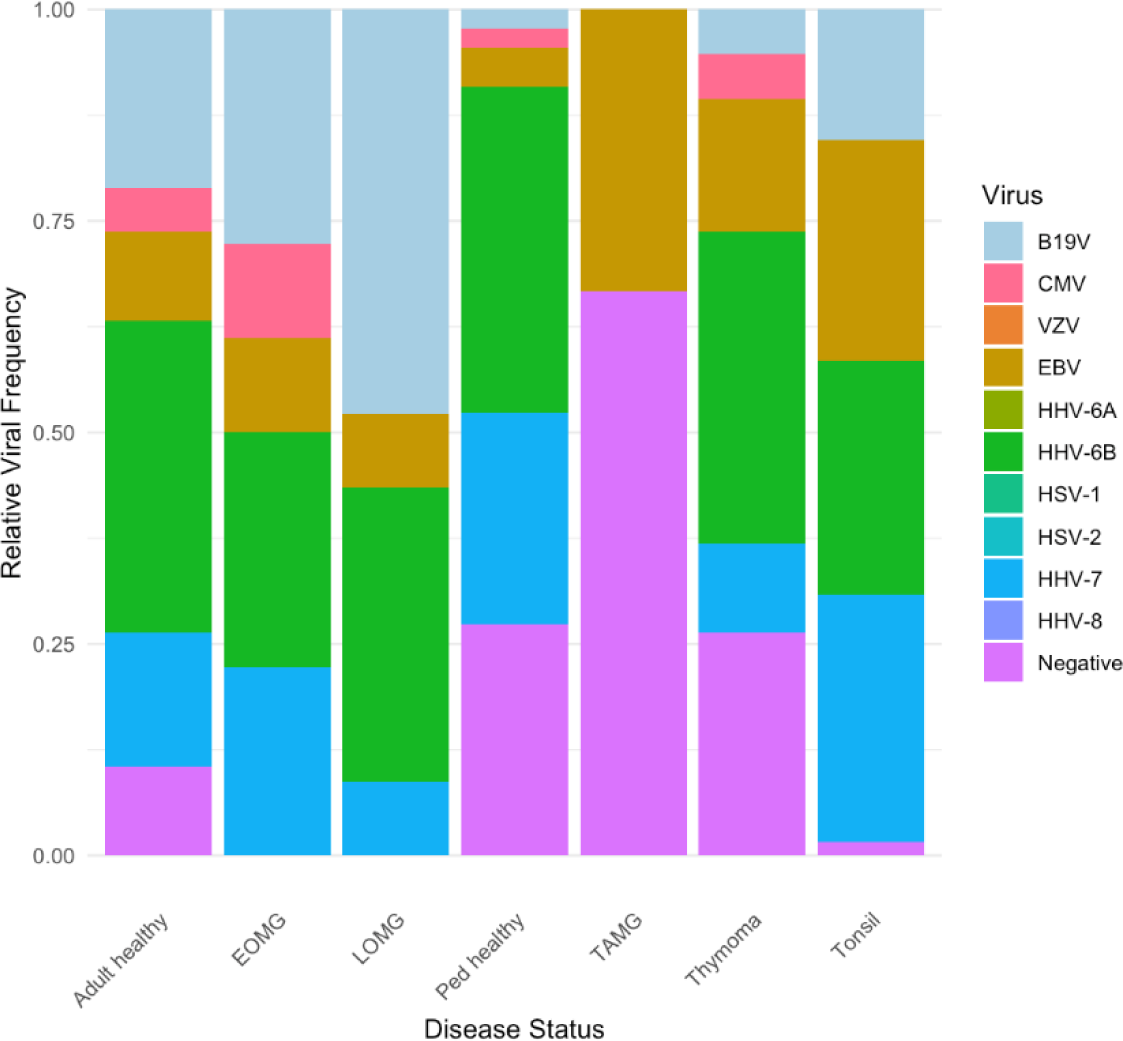
Comparative analysis of DNA-virus presence in subsets of MG patient thymic tissues and controls. Stacked bar plot illustrating the relative frequency of viral detection in each tissue group. The ’Negative’ group represents tissues PCR-Negative for all screened DNA viruses (herpesviruses and B19V).

